# REGN-COV2 antibody cocktail prevents and treats SARS-CoV-2 infection in rhesus macaques and hamsters

**DOI:** 10.1101/2020.08.02.233320

**Authors:** Alina Baum, Richard Copin, Dharani Ajithdoss, Anbo Zhou, Kathryn Lanza, Nicole Negron, Min Ni, Yi Wei, Gurinder S. Atwal, Adelekan Oyejide, Yenny Goez-Gazi, John Dutton, Elizabeth Clemmons, Hilary M. Staples, Carmen Bartley, Benjamin Klaffke, Kendra Alfson, Michal Gazi, Olga Gonzales, Edward Dick, Ricardo Carrion, Laurent Pessaint, Maciel Porto, Anthony Cook, Renita Brown, Vaneesha Ali, Jack Greenhouse, Tammy Taylor, Hanne Andersen, Mark G. Lewis, Neil Stahl, Andrew J. Murphy, George D. Yancopoulos, Christos A. Kyratsous

## Abstract

An urgent global quest for effective therapies to prevent and treat COVID-19 disease is ongoing. We previously described REGN-COV2, a cocktail of two potent neutralizing antibodies (REGN10987+REGN10933) targeting non-overlapping epitopes on the SARS-CoV-2 spike protein. In this report, we evaluate the in vivo efficacy of this antibody cocktail in both rhesus macaques and golden hamsters and demonstrate that REGN-COV-2 can greatly reduce virus load in lower and upper airway and decrease virus induced pathological sequalae when administered prophylactically or therapeutically. Our results provide evidence of the therapeutic potential of this antibody cocktail.

## Introduction

Fully human monoclonal antibodies are a promising class of therapeutics against SARS-CoV-2 infection (Cohen, 2020). To date, multiple studies have described discovery and characterization of potent neutralizing monoclonal antibodies targeting the spike glycoprotein of SARS-CoV-2 (Baum et al., 2020; Cao et al., 2020; Hansen et al., 2020; Ju et al., 2020; Liu et al., 2020; Pinto et al., 2020; Robbiani et al., 2020; Wang et al., 2020; Zost et al., 2020). However, evaluation of the efficacy of these antibodies in vivo is only beginning to emerge, and has largely focused on the prophylactic setting (Liu et al., 2020; Shi et al., 2020; Zost et al., 2020). Furthermore, as the animal models of SARS-CoV-2 infection and COVID-19 disease are still being developed, no single model has emerged as being more relevant for human disease. Indeed, based on the extremely diverse manifestations of COVID-19 in humans, multiple animal models may be needed to mimic various settings of human infection. The rhesus macaque model is widely used to assess efficacy of therapeutics and vaccines and displays a transient and mild course of the disease (Chandrashekar et al., 2020; Corbett et al., 2020; Deng et al., 2020; Mercado et al., 2020; Munster et al., 2020; Shan et al., 2020; van Doremalen et al., 2020; Yu et al., 2020). On the contrary, the golden hamster model manifests a much more severe form of the disease, accompanied by rapid weight loss and severe lung pathology (Imai et al., 2020; Rogers et al., 2020; Sia et al., 2020).

We previously described a cocktail of two fully human antibodies, REGN10933 and REGN10987, that bind to spike protein, potently neutralize SARS-CoV-2 and were selected as components of anti-viral antibody cocktail (REGN-COV2) to safeguard against mutational virus escape (Baum et al., 2020; Hansen et al., 2020). In this study, we utilized two different animal models, rhesus macaque and golden hamster, that capture the diverse pathology of SARS-CoV-2 infection and evaluated the in vivo efficacy of this antibody cocktail when used prophylactically or therapeutically. This assessment allows us to compare performance of the antibodies in diverse disease settings to more comprehensively understand the mechanisms by which monoclonal antibody therapies may limit viral load and pathology in infected individuals.

## Results

To evaluate the ability of REGN-COV2 to protect rhesus macaques from SARS-CoV-2 infection we initially assessed the impact of antibody administration prior to virus challenge (NHP Study #1). Animals were dosed with 50mg/kg of REGN-COV2 (25mg/kg of each antibody) through intravenous administration and challenged with 1×10^5 PFU of virus through intranasal and intratracheal routes 3 days post mAb dosing. Due to the relatively transient nature of the SARS-CoV-2 infection in rhesus macaques, the in-life portion of the study was limited to 5 days. To determine the impact of mAb prophylaxis on viral load in upper and lower airways we collected nasopharyngeal swabs on a daily basis and bronchoalveolar lavage (BAL) fluid on days 1, 3, and 5 post-challenge (Figure 1A). Both genomic and subgenomic RNA were measured to assess the impact of mAb prophylaxis on the dynamics of viral replication; while genomic RNA (gRNA) may reflect remaining viral inoculum as well as newly replicating virus, subgenomic RNA (sgRNA) should only result from newly replicating virus. For placebo-treated animals, the kinetics of viral load measures was as previously reported, with peak in viral load on day 2 post-challenge, although the majority of animals were still positive for viral RNA in nasal swabs on day 5; while the kinetics of gRNA and sgRNA were similar, sgRNA levels were about a hundred-fold lower, consistent with what others have reported (Chandrashekar et al., 2020; Mercado et al., 2020; Yu et al., 2020; Zost et al., 2020). For animals receiving REGN-COV2 prophylaxis we observed markedly accelerated clearance of gRNA with almost complete ablation of sgRNA in the majority of the animals, showing that REGN-COV2 can almost completely block establishment of virus infection; this pattern was observed across all measurements in both nasopharyngeal swabs and BAL compared to placebo animals, demonstrating that mAbs administered prophylactically can greatly reduce viral load in both upper and lower airways (Figure 1B).

**Figure 1.**
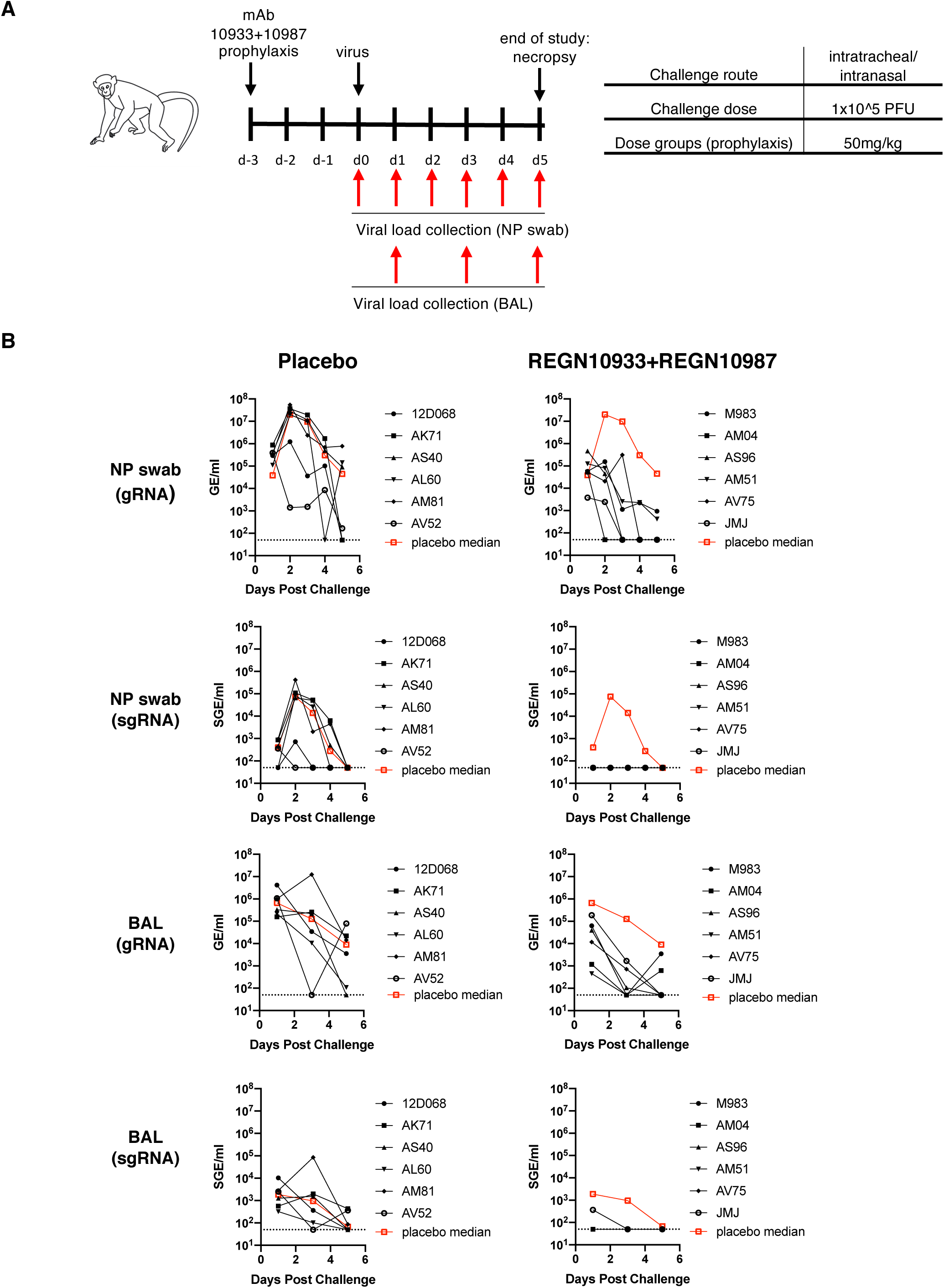
Prophylactic efficacy of REGN-COV2 in the rhesus macaque model of SARS-CoV-2 infection (NHP Study #1) **(A)** Overview of study design. (**B)** Impact of REGN-COV2 prophylaxis on viral genomic RNA (gRNA) and subgenomic RNA (sgRNA) in nasopharyngeal swabs and bronchioalveolar lavage (BAL) fluid.

A second prophylaxis study (NHP Study #2) was designed to test whether REGN-COV2 could protect against a 10-fold higher viral inoculum (1.05×10^6 PFU), and compared the 50mg/kg dose of REGN-COV2 (25mg/kg of each antibody) with a much lower dose (Figure 2A). Nasopharyngeal and oral swabs were collected and used to measure virus genomic and subgenomic virus RNA. We observed that 50mg/kg of REGN-COV2 administered 3 days prior to virus challenge was once again able to minimize virus replication even when animals were challenged with this 10-fold higher viral challenge (Figure 2B), while the prophylactic effect was greatly diminished with the 0.3mg/kg dose. Interestingly, in this study we observed increased impact of mAb treatment on viral load in oral swabs versus nasopharyngeal swabs, potentially indicating that mAb treatment may impact multiple physiological sources of virus replication differentially. Additional studies in animal models and humans will be needed to assess whether this is really the case.

**Figure 2.**
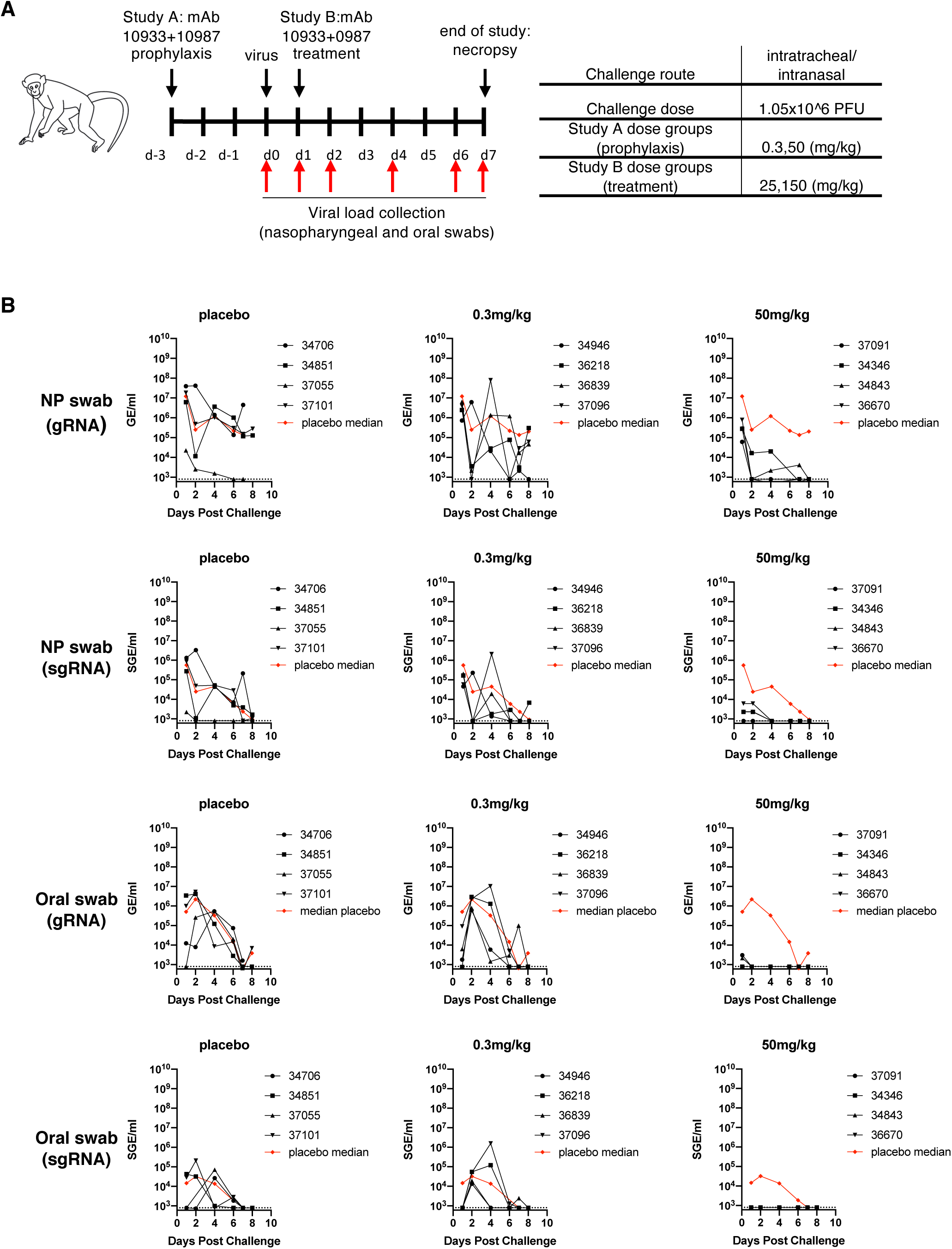
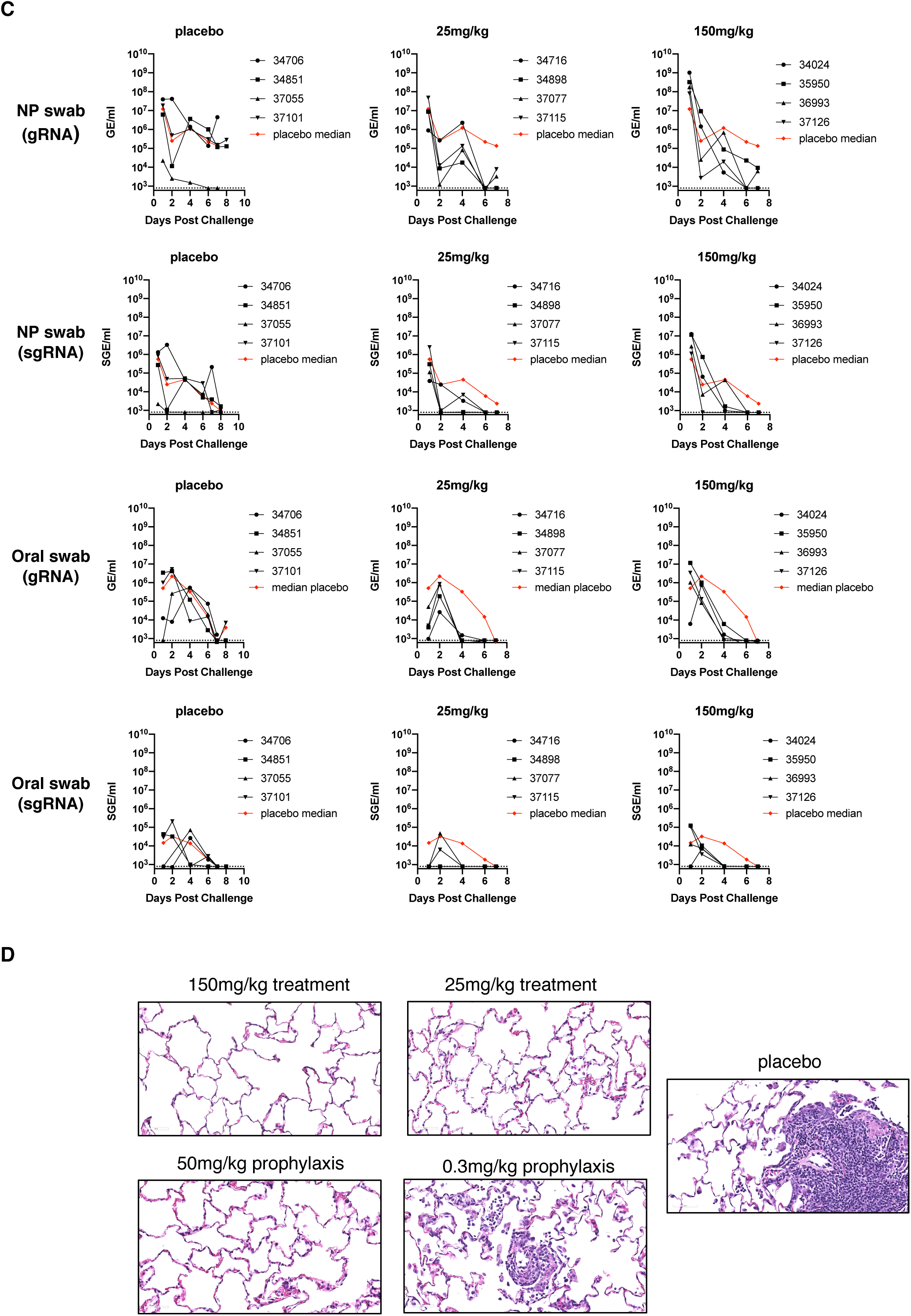
Prophylactic and therapeutic efficacy of REGN-COV2 in the rhesus macaque model of SARS-CoV-2 infection (NHP Study #2) **(A)** Overview of study design. (**B)** Impact of REGN-COV2 prophylaxis on viral genomic RNA (gRNA) and subgenomic RNA (sgRNA) in nasopharyngeal swabs and oral swabs. (**C)** Impact of REGN-COV2 treatment on viral genomic RNA (gRNA) and subgenomic RNA (sgRNA) in nasopharyngeal swabs and oral swabs. (**D)** representative images of histopathology in lungs of treated and placebo animals.

Next, we assessed the impact of REGN-COV2 in the treatment setting by dosing animals challenged with the higher 1×10^6 PFU of SARS-CoV-2 virus at 1-day post-infection with 25mg/kg or 150mg/kg of the antibody cocktail (Figure 2A). By day 1 post-challenge the animals already reached peak viral load as measured by both genomic and subgenomic RNA, mimicking a likely early treatment clinical scenario of COVID-19 disease, since it has been shown that most SARS-CoV-2 infected individuals reach peak viral loads relatively early in the disease course and often prior or just at start of symptom onset (He et al., 2020; Zou et al., 2020). Compared to placebo treated animals, REGN-COV2 treated animals displayed accelerated viral clearance in both nasopharyngeal and oral swabs samples, including both genomic and subgenomic RNA samples (Figure 2C), clearly demonstrating that the monoclonal antibody cocktail can impact virus load even when administered post infection. Similar to the prophylaxis study, the decrease in viral load appeared more dramatic in oral swabs versus nasopharyngeal swabs. Both treatment groups displayed similar kinetics of virus clearance, suggesting that 25mg/kg and 150mg/kg demonstrate similar efficacy in this study. The treated animals in the 150mg/kg group displayed approximately 10-fold higher titers on day 1, at the time of mAb administration, therefore potentially masking enhanced effect of a higher drug dose. Similar impact of mAb treatment was observed on genomic and subgenomic RNA for both NP and oral samples, indicating the mAb treatment is directly limiting viral replication in these animals (Figure 2C).

The two antibody components of REGN-COV2 were selected to target non-overlapping sites on the spike protein to prevent selection of escape mutants, which were readily detectable with single mAb treatment (Baum et al., 2020). To assess whether any signs of putative escape mutants are observed in an in vivo setting with authentic SARS-CoV-2 virus, we performed RNAseq analysis on all RNA samples obtained from all animals from the study. Analysis of the spike protein sequence identified mutations in NHP samples that were not present in the inoculum virus (Figure S1) further indicating that the virus is actively replicating in these animals. However, we did not observe any mutations that were unique to treated animals; all identified mutations were either present in the inoculum or in both treated and placebo animals, indicating that they were likely selected as part of virus replication in NHPs and were not selected by mAb treatment.

We next performed pathology analyses of lungs of infected animals. All four placebo monkeys showed evidence of lung injury characterized in three monkeys by interstitial pneumonia (Figure 2D), with minimal to mild infiltration of mononuclear cells (lymphocytes and macrophages) in the septa, perivascular space, and/or pleura. In these three animals, the distribution of lesions was multifocal and involved 2-3 of the 4 lung lobes. Accompanying these changes were alveolar infiltration of lymphocytes, increased alveolar macrophages, and syncytial cells. Type II pneumocyte hyperplasia was also observed in occasional alveoli. In the fourth placebo monkey, lung injury was limited to type II pneumocyte hyperplasia, suggestive of a reparative process secondary to type I pneumocyte injury. Overall, the histological lesions observed in the placebo animals were consistent with an acute SARS-CoV-2 infection. In the prophylactic groups, 3 of 4 animals in the low dose (0.3mg/kg) and 1 of 4 animals in the high dose (50mg/kg) groups showed evidence of interstitial pneumonia (Table S1) that was generally minimal and with fewer histological features when compared to the placebo group. In the one affected high dose group animal, only 1 of the 4 lung lobes had a minimal lesion. In the therapeutic treatment groups, 2 of 4 low dose (25mg/kg) and 2 of 4 high dose (150mg/kg) treated animals showed evidence of interstitial pneumonia. In all affected low and high dose animals, only 1 of 4 lung lobes had lesions. Finally, there was no test article related toxicities observed at any of the doses tested. In summary, the incidence of interstitial pneumonia (number of animals as well as number of lung lobes affected) and the severity were reduced in both prophylactic and therapeutic treatment modalities, compared to placebo. The analyses demonstrated that prophylactic and therapeutic administration of REGN-COV2 greatly reduced virus induced pathology in rhesus macaques and showed a clean safety profile.

Unlike rhesus macaques which present with a mild clinical course of disease and transient virus replication when infected with SARS-CoV-2, which may mimic mild human disease, the golden hamster model is more severe, with animals demonstrating readily observable clinical disease, including rapid weight loss accompanied by very high viral load in lungs, as well as severe lung pathology. Thus, this model may more closely mimic more severe disease in humans, although more extensive characterization of this model and severe human disease is needed to better understand similarities and differences in pathology. To evaluate the ability of REGN-COV2 to alter the disease course in this model, we designed a study which evaluated the prophylactic and treatment efficacy of the antibodies (Figure 3A). Administration of 50, 5 or 0.5mg/kg of REGN-COV2 2 days before challenge with 2.3×10^4 PFU dose of SARS-CoV-2 virus resulted in dramatic protection from weight loss at all doses. This protection was accompanied by greatly decreased viral load in the lungs at the end of the study (day 7 post infection) (Figure 3C). Interestingly we did observe high gRNA and sgRNA levels in the lungs of a few treated animals, however these individual animals did not show decreased protection from weight loss than the animals with much lower viral loads. It is possible that mAb treatment may provide additional therapeutic benefit in this model not directly associated with viral load decrease. Alternatively, it is possible that the increased detected viral RNA may not necessarily be associated with infectious virus. As viral replication and lung pathology in the hamster model occur very rapidly, the treatment setting represents a high bar for demonstrating therapeutic efficacy. We were able to observe therapeutic benefit in animals treated with 50mg/kg and 5mg/kg doses of REGN-COV2 combination 1-day post viral challenge (Figure 3B). Taken together the two hamster studies clearly demonstrate that REGN-COV2 can alter the course of infection in the hamster model of SARS-COV-2 either when administered prophylactically or therapeutically.

**Figure 3.**
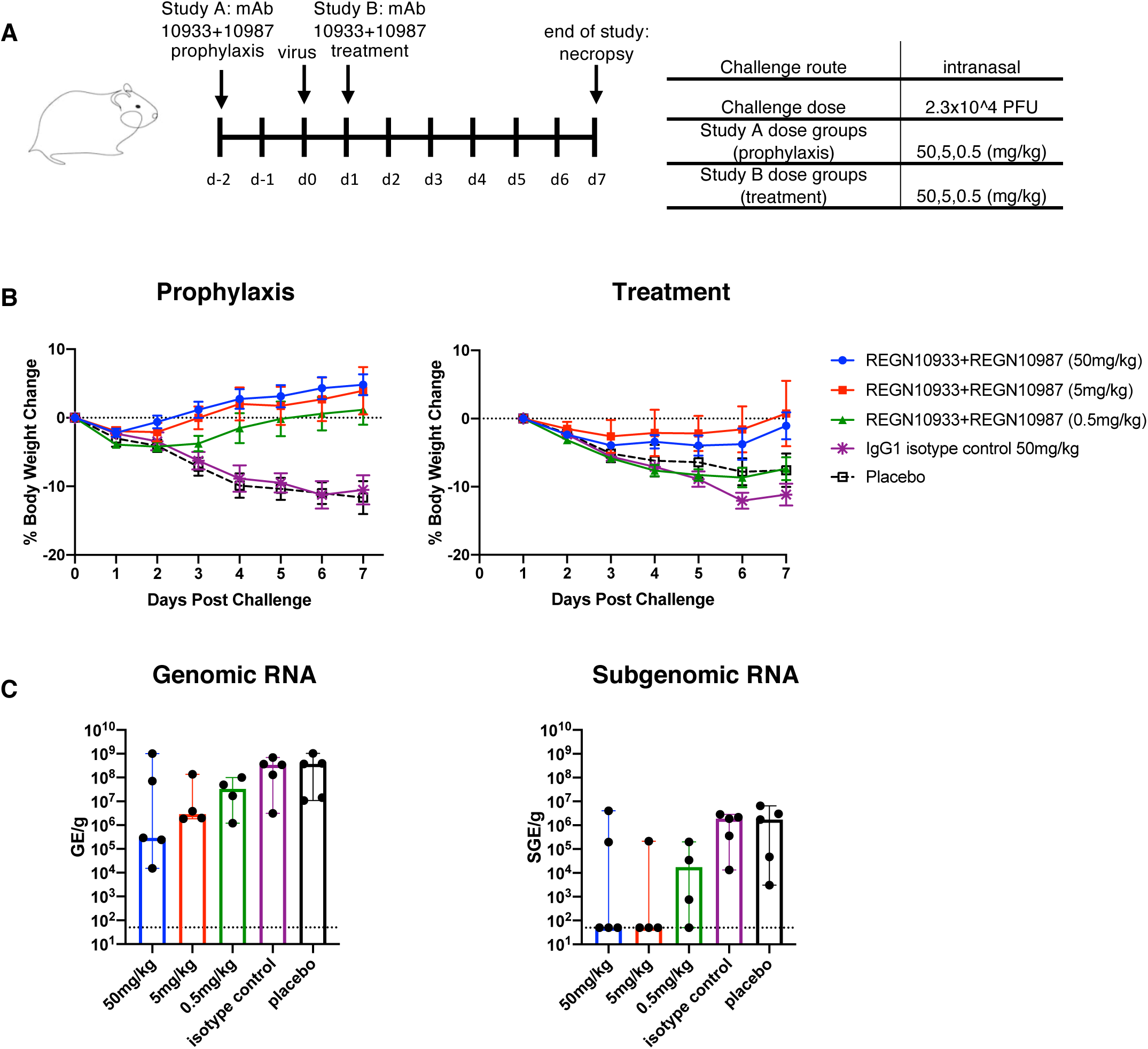
Efficacy of REGN-COV2 in treatment and prophylaxis in the golden Syrian hamster model of SARS-CoV-2 infection. **(A)** Study design overview, **(B)** Impact of REGN-COV2 on weight loss in prophylaxis and treatment, (**C)** Impact of REGN-COV-2 prophylaxis on levels of gRNA and sgRNA in hamster lungs (7dpi).

## Discussion

In this study, we assessed the in vivo prophylactic and treatment efficacy of the REGN-COV2 mAb cocktail in two animal models, one of mild disease in rhesus macaques and one of severe disease in golden hamsters. Our results demonstrated that the antibodies are efficacious in both animal models, as measured by reduced viral load in the upper and lower airways, reduced virus induced pathology in the rhesus macaque model, and by limited weight loss in the hamster model.

The ability of REGN-COV2 to almost completely block detection of subgenomic species of SARS-COV-2 RNA matches or exceeds the effects recently shown in vaccine efficacy studies using the same animal models (Corbett et al., 2020; Gao et al., 2020; Mercado et al., 2020; Patel A. et al., 2020; van Doremalen et al., 2020). Additionally, the observed accelerated reduction of upper airway virus load in rhesus macaques treated with REGN-COV2 contrasts the lack of impact on viral load in remdesivir treated animals, where reduced viral load could only be observed in lower airways with no differences in nasal viral RNA levels (Williamson et al., 2020). These findings highlight the therapeutic potential of REGN-COV2 to both protect from and treat SARS-COV-2 disease. Additionally, the impact of REGN-COV2 prophylaxis on viral RNA levels in nasopharyngeal and oral swabs may indicate the potential to not only prevent disease in the exposed individual but also to limit transmission.

To our knowledge this is the first report demonstrating ability of any therapeutic to limit weight loss in the treatment setting of SARS-CoV-2 infection in the hamster model, indicating potential benefit of antibody treatment in the context of a severe infection. Further understanding of both the hamster and the macaque model and how their disease course and pathology mimics the breadth of human COVID-19 disease may help to gain more in depth understanding of how mAb therapeutics may confer clinical benefit.

Importantly, in our studies we did not observe any signs of increased viral load and/or worsening of pathology in presence of antibodies at either high or low doses in either animal model. Potential for antibody mediated enhancement of disease (ADE) is a serious concern for antibody-based therapeutics and vaccines. And although a recent report showed ability of some anti-spike mAbs to mediate pseudovirus entry into FcγR expressing cell lines, these data do not address whether similar behavior would be observed with authentic SARS-CoV-2 virus and primary immune cells (Wang S. et al., 2020). Our results are consistent with no evidence of enhanced disease in clinical studies assessing convalescent plasma therapy (Li et al., 2020).

In conclusion, our data provide evidence that REGN-COV2 based therapy may offer clinical benefit in both prevention and treatment settings of COVID-19 disease, where it is currently being evaluated (clinicaltrials.gov NCT04426695, NCT04425629 and NCT 04452318).

## Methods

### Studies conducted at BIOQUAL (NHP Study #1 and Hamster Study)

#### Ethics Statement and Animal Exposure

Animal research was conducted under BIOQUAL Institute Institutional Animal Care and Use Committee (IACUC)-approved protocols, 20-070P (hamster study) and 20-069P (NHP study) in compliance with the Animal Welfare Act and other federal statutes and regulations relating to animals and experiments involving animals. BIOQUAL is accredited by the Association for Assessment and Accreditation of Laboratory Animal Care International and adheres to principles stated in the Guide for the Care and Use of Laboratory Animals, National Research Council. Animals were monitored at least twice daily, and enrichment included commercial toys and food supplements. Prior to all blood collections, animals were anesthetized using Ketamine (NHPs) or Ketamine/Xylazine (hamsters). At the end of the study, animals were euthanized with an intravenous (NHPs) or intraperitoneal (hamsters) overdose of sodium pentobarbital.

#### Rhesus macaque study

A total of 12 naïve rhesus macaques of Indian origin (purpose bred, *Macaca mulatta*) were used in the study. Animals were distributed to treatment groups based on age distribution. Antibodies or saline were administered through intravenous infusion. Animals were challenged with 1.1×10^5 PFU (USA-WA1/2020 (NR-52281; BEI Resources) total dose of virus divided between intranasal and intratracheal routes. Virus was administered using a 3mL syringe to drop-wise instill 1 mL by the intranasal (IN) route (0.5 mL in each nare) and using a French rubber tube, administer 1 mL via the intratracheal (IT) route. Viral titers were collected by nasal swabs (2x Copan flocked per animal, placed into one vial each with 1mL PBS) and bronchioalveolar lavage (BAL) using 10 mL saline via a rubber feeding tube. Collected swabs and BAL aliquots were stored at −80°C until viral load analysis.

#### Hamster study

A total of 50 golden hamsters, male and female, 6-8 weeks old were used in the study. Animals were weighed prior to the start of the study. The animals were monitored twice daily for signs of COVID-19 disease (ruffled fur, hunched posture, labored breathing, a.o.) during the study period. Body weights were measured once daily during the study period. Antibodies were dosed through intraperitoneal (IP) injection. Animals were challenged with 2.3×10^4 PFU of (USA-WA1/2020 (NR-52281; BEI Resources) by administration of 0.05mL of viral inoculum dropwise into each nostril. Tissues were sampled for viral load assays by collecting two small pieces (0.1-0.2 gram each) from the lung (total of 4 pieces, 2 per tissue). Tissues were stored at −80°C until viral load analysis.

#### Cells and Virus

Vero E6 cells (ATCC, catalog number CRL 1586) were grown in Dulbecco’s modified essential media (DMEM; Gibco) with 10% heat-inactivated fetal bovine serum (FBS; Gibco) at 37°C with 5% CO2. SARS-CoV-2 (P4) isolate USA-WA1/2020 (BEI resources NR-52281, GenBank accession number MN985325.1) was used to generate the animal exposure stock (P5). The stock was generated by infecting Vero E6 cells at an MOI of 0.002 in DMEM containing 2% FBS; viral supernatant was harvested at four days post infection. The stock has been confirmed to be SARS-CoV-2 via deep sequencing and confirmed to be free of adventitious agents. The viral titer was determined to be 2.3 ×10^5 PFU/mL.

#### Quantitative RT-PCR Assay for SARS-CoV-2 RNA

The amounts of RNA copies per mL bodily fluid or per gram tissue were determined using a qRT-PCR assay. The qRT-PCR assay utilized primers and a probe specifically designed to amplify and bind to a conserved region of nucleocapsid gene of coronavirus. The signal was compared to a known standard curve and calculated to give copies per mL. For the qRT-PCR assay, viral RNA was first isolated from nasal wash using the Qiagen MinElute virus spin kit (cat. no. 57704). For tissues it was extracted with RNA-STAT 60 (Tel-test”B”)/ chloroform, precipitated and resuspended in RNAse-free water. To generate a control for the amplification reaction, RNA was isolated from the applicable SARS-CoV-2 stock using the same procedure. qPCR assay was performed with Applied Biosystems 7500 Sequence detector and amplified using the following program: 48°C for 30 minutes, 95°C for 10 minutes followed by 40 cycles of 95°C for 15 seconds, and 1 minute at 55°C. The number of copies of RNA per mL was calculated by extrapolation from the standard curve and multiplying by the reciprocal of 0.2 mL extraction volume. This gives a practical range of 50 to 5 × 10^8^ RNA copies per mL for nasal washes or per gram of tissue.

Primers/probe sequences:

2019-nCoV_N1-F: 5’-GAC CCC AAA ATC AGC GAA AT-3’

2019-nCoV_N1-R: 5’-TCT GGT TAC TGC CAG TTG AAT CTG-3’

2019-nCoV_N1-P: 5’-FAM-ACC CCG CAT TAC GTT TGG TGG ACC-BHQ1-3’

#### Quantitative RT-PCR Assay for SARS-CoV-2 subgenomic RNA

SARS-CoV-2 E gene subgenomic mRNA (sgRNA or sgmRNA) was assessed by RT-PCR using primers and probes as previously described(Chandrashekar et al., 2020). Briefly, to generate a standard curve, the SARS-CoV-2E gene sgRNA was cloned into a pcDNA3.1 expression plasmid; this insert was transcribed using an AmpliCap-Max T7 High Yield MessageMaker Kit (Cellscript) to obtain RNA for standards. Prior to RT-PCR, samples collected from challenged animals or standards were reverse-transcribed using Superscript III VILO (Invitrogen) according to the manufacturer’s instructions. A Taqman custom gene expression assay (ThermoFisher Scientific) was designed using the sequences targeting the E gene sgRNA20. Reactions were carried out on a QuantStudio 6 and 7 Flex Real-Time PCR System (Applied Biosystems) according to the manufacturer’s specifications. Standard curves were used to calculate sgRNA in copies per ml or per swab; the quantitative assay sensitivity was 50 copies per ml or per swab. This gives a practical range of 50 to 5 × 10^7 RNA copies per mL for nasal washes, and for tissues the viral loads are given per gram.

Subgenomic RNA Primers:

SG-F: CGATCTTGTAGATCTGTTCCTCAAACGAAC

SG-R: ATATTGCAGCAGTACGCACACACA

PROBE: FAM-ACACTAGCCATCCTTACTGCGCTTCG-BHQ

### Studies conducted at Texas Biomedical Research Institute (NHP Study #2)

#### Ethics Statement and Nonhuman Primate Exposure

Animal research was conducted under Texas Biomedical Research Institute Institutional Animal Care and Use Committee (IACUC)-approved protocol (1721MM) in compliance with the Animal Welfare Act and other federal statutes and regulations relating to animals and experiments involving animals. Texas Biomedical Research Institute is accredited by the Association for Assessment and Accreditation of Laboratory Animal Care International and adheres to principles stated in the Guide for the Care and Use of Laboratory Animals, National Research Council. Animals were monitored at least twice daily and enrichment included commercial toys and food supplements. Prior to all blood collections, animals were anesthetized using Telazol (Zoetis Inc., Parsippany-Troy Hills, NJ, USA). At the end of the study, animals were euthanized with an intravenous overdose of sodium pentobarbital.

#### Animal challenge

Twenty-four (24) rhesus macaques (13 female and 11 males) were used in this study, and randomly assigned to one of six groups. Animals were obtained from the Southwest National Primate Research Center (SNPRC) colony and were between 2.5 and 6 years of age and approximately 3 to 10 kg at the time of study enrollment. On Study Day 0, each NHP was exposed at ABSL-4 with a targeted dose of 1.05 × 10^6^ PFU of SARS-CoV-2 in a total volume of 500 µl (5.25 × 10^5^ PFU in 250 µl via intranasal route and 5.25 × 10^5^ PFU in 250 µl via intratracheal route). Intranasal delivery was via a mucosal atomization device (Teleflex Intranasal Mucosal Atomization Device LMA MAD Nasal Device), which allows for IN delivery of atomized particles 30 - 100 microns in size, which model droplet transmission. Mucosal atomization devices have been developed for safe and efficient drug delivery to administer drugs that are United States Food and Drug Administration (U.S. FDA) approved for IN delivery. Intratracheal delivery used a Tracheal Mucosal Atomization Device (Teleflex Laryngo-Tracheal Mucosal Atomization Device LMA MADGIC). Animals were exposed in ascending order based on Texas Biomed animal ID in order to minimize timing bias for treatment administration. On Day −3 relative to exposure, prophylactic group animals were sedated and received treatment. On Day 1 (post virus exposure), therapeutic group animals were sedated and received treatment. Treatment was administered via intravenous injection over the course of approximately 90 seconds.

#### Cells and Virus

Vero E6 cells (VERO C1008, catalog number NR-596, BEI resources) were grown in Dulbecco’s modified essential media (DMEM; Gibco) with 10% heat-inactivated fetal bovine serum (FBS; Gibco) at 37°C with 5% CO2. SARS-CoV-2 isolate USA-WA1/2020 (BEI resources NR-52281, GenBank accession number MN985325.1) was used to generate the animal exposure stock. A fourth cell-culture passage (P4) of SARS-CoV-2 was obtained from in 2020 and propagated at Texas Biomedical Research Institute. The fourth cell-culture passage (P4) stock virus obtained from BEI was passaged one time to generate a master stock by infecting Vero E6 cells at a multiplicity of infection (MOI) of approximately 0.001 in DMEM containing 2% FBS; viral supernatant was harvested at 3 days post infection. The P5 stock was used to generate the exposure stock by infecting Vero E6 cells at an MOI of 0.02 in DMEM containing 2% FBS; viral supernatant was harvested at three days post infection. The stock has been confirmed to be SARS-CoV-2 via deep sequencing and confirmed to be free of adventitious agents. The viral titer was determined to be 2.1 × 10^6^ PFU/mL.

#### RNA extraction for viral load determination via RT-qPCR

Samples were inactivated using TRIzol LS Isolation Reagent (Invitrogen): 250 µL of test sample were mixed with 750 µL TRIzol LS. Inactivation controls were prepared with each batch of samples. Prior to extraction, 1 × 10^3^ pfu of MS2 phage (Escherichia coli bacteriophage MS2, ATCC) was added to each sample to assess extraction efficiency RNA extraction was performed using the EpMotion M5073c Liquid Handler (Eppendorf) and the NucleoMag Pathogen kit (Macherey-Nagel). Extraction controls were prepared with each batch of samples. After processing, the presence of the eluate was confirmed and the extracted RNA was stored at - 80°C±10°C.

#### Determination of Viral load via RT-qPCR

5 µL RNA sample was taken to duplex RT-qPCR reaction detecting both SARS-CoV-2 and MS2 phage. Two assays were used to assess SARS-CoV-2 present in the samples. The CDC-developed 2019-nCoV_N1 assay was used to target a region of the N gene. SARS-CoV-2_N1 probe (ACCCCGCATTACGTTTGGTGGACC) is labeled with 6-FAM fluorescent dye. The forward primer sequence is: GACCCCAAAATCAGCGAAAT, and the reverse primer sequence is: TCTGGTTACTGCCAGTTGAATCTG. A secondary qPCR assay to measure subgenomic RNA was also performed to target a region of E (Envelope)(Corman et al., 2020; Wolfel et al., 2020) The probe is also labeled with 6-FAM fluorescent dye (ACACTAGCCATCCTTACTGCGC TTCG). The forward primer sequence is: CGATCTCTTGTAGATCTGTTCTC, and the reverse primer sequence is: ATATTGCAGCAGTACGCACACA. The MS2 probe is labeled with VIC fluorescent dye. Both assays used the TaqPath™ 1-Step RT-qPCR Master Mix, CG (ThermoFisher) and were performed on a QuantStudio 3 instrument (Applied Biosystems). QuantStudio Design and Analysis Software (Applied Biosystems) was used to run and analyze the results. Cycling parameters were set as follows: Hold stage 2 min at 25°C, 15 min at 50°C, 2 min at 95°C. PCR stage: 45 cycles (N1 assay) or 40 cycles (E assay) of 3 sec at 95°C, 30 sec at 60°C. The average Ct value for MS2 phage was calculated for all processed samples and SARS-CoV-2 quantification only performed in samples in which the MS2 Ct value was lower than Average MS2 + 5%.

#### Histopathology

Necropsies were conducted by BSL-4 personnel in accordance with SOP Texas Biomed 916 and selected tissue samples (tracheobronchial lymph node, nasal cavity, trachea, heart, liver, spleen, kidney, and all 4 right lung lobes) were collected. Tissues were fixed by immersion in 10% neutral-buffered formalin for a minimum of fourteen days, then trimmed, routinely processed, and embedded in paraffin. Sections of the paraffin-embedded tissues were cut at 5 µm thick, and histology slides were deparaffinized, stained with hematoxylin and eosin (H&E), cover slipped, and labeled. Slides were blindly evaluated by a board-certified veterinary pathologist.

#### Virus RNA Sequencing

10 ul of RNA combined with 25 ng Human Universal Reference RNA (Agilent) was purified by PureBeads (Roche Sequencing). cDNA synthesis was performed using SuperScript™ IV First-Strand Synthesis System (Thermal Fisher) following vendor’s protocol. Then one half of cDNA (10 ul) was used to generate libraries using Swift Normalase™ Amplicon Panel (SNAP) SARS CoV-2 Panel (Swift Biosciences) following vendor’s protocol. Sequencing was run on NextSeq (Illumina) by multiplexed paired-read run with 2×150 cycles.

#### RNAseq data analysis

RNAseq analysis was perform using Array Studio software package platform (Omicsoft). Quality of paired-end RNA Illumina reads was assessed using the “raw data QC of RNA-Seq data suite”. Minimum and maximum read length, total nucleotide number, and GC% were calculated. Overall quality report was generated summarizing the quality of all reads in each sample, along each base pair. Swift amplicon bulk RNA-seq reads were aligned to the SARS-COV-2 reference genome Wuhan-Hu-1 (MN908947) using Omicsoft Sequence Aligner (OSA) version 4. The alignments were sorted by read name, and primers were clipped by the complementary Swiftbiosciences primerclip software (v0.3.8) (https://github.com/swiftbiosciences/primerclip). Reads were trimmed by quality score using default parameters (when aligner encountered nucleotide in the read with a quality score of 2 or less, it trimmed the remainder of the read). OSA outputs were analyzed and annotated using Summarize Variant Data and Annotate Variant Data packages (Omicsoft). The rest of the analysis focused on the genome section encoding the Spike protein. Using custom scripts, target coverage was summarized for each sample and SNPs calling was calculated. The frequency of viral mutations inferred from the sequencing reads were calculated if mutated reads were higher than 1% relative to total number reads.

## Acknowledgments

The following reagent was deposited by the Centers for Disease Control and Prevention and obtained through BEI Resources, NIAID, NIH: SARS-Related Coronavirus 2, Isolate USA-WA1/2020, NR-52281.

## Funding

A portion of this project has been funded in whole or in part with Federal funds from the Department of Health and Human Services; Office of the Assistant Secretary for Preparedness and Response; Biomedical Advanced Research and Development Authority, under OT number: HHSO100201700020C.

## Author contributions

A.B., N.S, A.J.M, G.D.Y., C.A.K. conceptualized and designed experiments. Y.G.G, J.D., E.C., H.S., C.B., B.K., O.G., E.D., L.P., M.P., A.C., R.B., V.A., J.G., T.T., performed experiments and A.B., R.C., D.A., A.O, K.A., R.C., M.G., H.A., M.G.L., M.A., G.D.Y., C.A.K. analyzed data. R.C., K.L., N.N., M.N., Y.W. prepared sequencing libraries and performed bioinformatics analysis A.B. and C.A.K. wrote the paper. C.A.K. acquired funding.

## Competing interests

Regeneron authors own options and/or stock of the company. This work has been described in one or more pending provisional patent applications. N.S, A.J.M., G.D.Y. and C.A.K. are officers of Regeneron.

---

**Figure S1.**
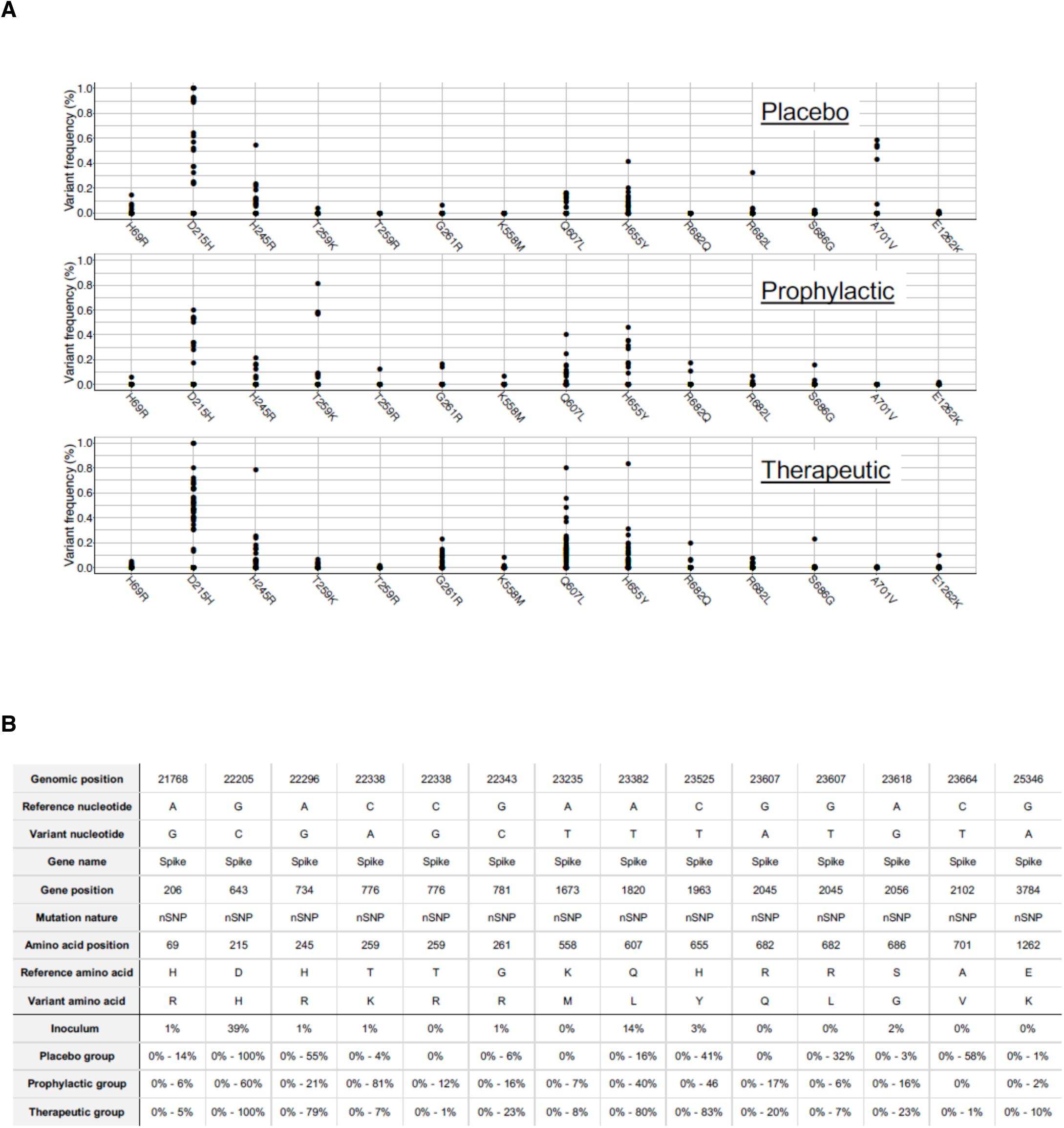
RNAseq analysis of viral RNA from NHP study #2. **(A)** Virus RNA was sequenced and RNAseq analysis was performed to identify amino acid changes relative to virus inoculum sequence. The graph shows the frequencies of all amino acid changes identified in the spike protein across all virus sequences. Each dot represents the frequency of the corresponding amino acid change in a specific virus sample. Samples are grouped based on treatment regiment: isotype control (Placebo), therapeutic antibodies administered prior (Prophylactic) or following (Treatment) viral challenge. (**B)** Detailed genomic information on all amino acid changes identified within the spike protein sequence across all samples. For each sample, the frequency of all mutations has been calculated. These frequencies are shown as percentage of the virus population with the amino acid change in the input virus or as range of frequency percentages (lowest to highest %) in the virus populations isolated from the placebo, prophylactic and therapeutic groups.

**Table S1.**
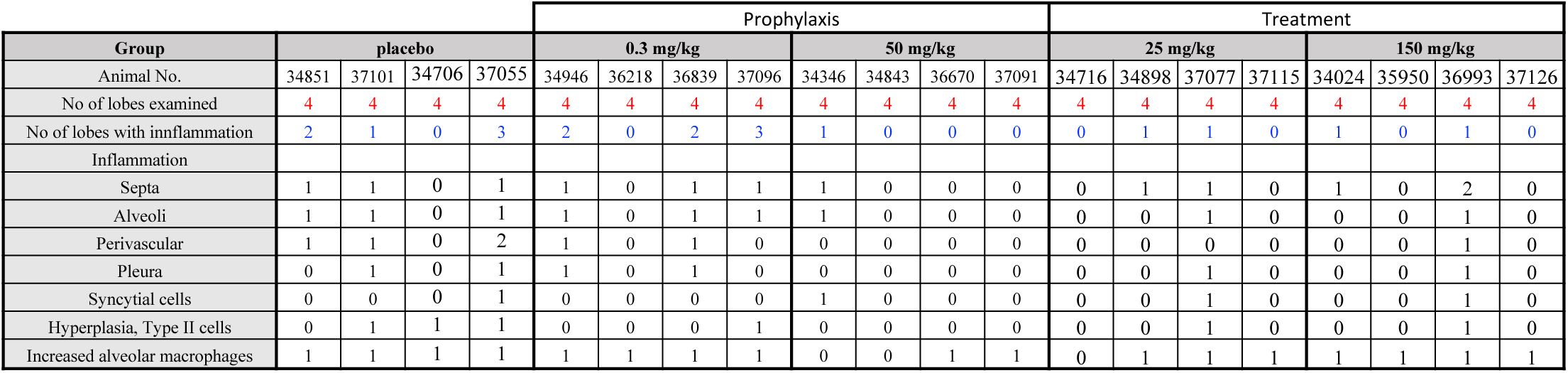
Pathology analysis in rhesus macaque lungs (NHP Study #2). Pathology scores in individual animals treated with either REGN-COV-2 or placebo. Severity score of lesions: Minimal (1); Mild (2); Moderate (3); Severe (4)

